# MAP-PRS: Multi-Ancestry Portfolio-Based Polygenic Risk Scores

**DOI:** 10.1101/2025.10.15.682516

**Authors:** Lokendra S. Thakur, Nilanchali Singh, Gurpreet Bharj

## Abstract

Polygenic Risk Scores (PRS) are emerging tools for predicting an individual’s genetic risk for complex diseases. However, their usefulness in clinical practice remains limited because most existing models are based on data from people of European ancestry, leading to reduced accuracy and stability in other populations. This imbalance restricts the equitable use of PRS in precision medicine. To overcome these limitations, we introduce **Multi-Ancestry Portfolio-Based Polygenic Risk Scores (MAP-PRS)**—a new framework that combines mathematical modeling and data science principles to improve both fairness and reliability in genetic risk prediction across populations. MAP-PRS treats each ancestry-specific PRS as part of a “portfolio,” similar to how investments are managed in finance, balancing two key aspects: *predictive return* (how well the score predicts disease) and *risk complexity* (how uncertain or ancestry-specific the prediction is). By jointly optimizing these factors, MAP-PRS identifies the best combination of ancestry-informed PRS models that maximize predictive accuracy while minimizing bias and instability. This approach also uses advanced computational tools—such as Bayesian modeling, machine learning, and generative neural networks—to refine risk estimates, incorporate environmental and lifestyle factors, and increase representation from under-studied populations. In doing so, MAP-PRS supports more inclusive, equitable, and interpretable precision medicine. As an initial demonstration, MAP-PRS has been applied to predict Type 2 Diabetes (T2D) risk in European ancestry populations, establishing a foundation for broader, multi-ancestry implementation. Future extensions will include additional diseases, such as cervical cancer and HPV susceptibility, endometrioid ovarian cancer, and Alzheimer’s disease—bringing us closer to clinically actionable and globally equitable genetic risk prediction.

## 1 Introduction

Polygenic Risk Scores (PRS) are a powerful tool for predicting disease risk but suffer from ancestry bias, limiting their effectiveness in genetically diverse populations such as India.^**1, 2**^ Existing PRS models are largely derived from European-centric GWAS datasets, leading to suboptimal risk predictions in non-European groups. Several PRS methodologies have been developed, each with different assumptions and limitations. These include: 1. Clumping and Thresholding (C+T) PRS – A traditional approach that selects SNPs based on linkage disequilibrium (LD) pruning and predefined significance thresholds.^**3**^ 2. LDpred – A Bayesian method that improves PRS accuracy by adjusting effect sizes based on linkage disequilibrium patterns.^**4**^ 3. PRS-CS – An extension of LDpred that uses continuous shrinkage priors to improve prediction, particularly in ancestrally diverse populations.^**5**^ 4. Bayesian Polygenic Risk Scores (BPRS) – Incorporates hierarchical modeling and Bayesian shrinkage to refine effect size estimates.^**6**^ 5. Machine Learning-Based PRS – Includes methods like LASSO regression, support vector machines (SVMs), and neural networks to capture complex interactions in genomic data.^**7**^ 6. Multi-Ancestry PRS – Combines PRS trained on multiple ancestry groups to improve transferability across populations.^**8**^ 7. Functional Annotation-Informed PRS – Integrates functional genomic annotations to prioritize causal variants for disease prediction.^**9**^

Despite these advancements, PRS models still exhibit limited generalizability across non-European populations. Therefore, there is a need for equitable and population-specific PRS frameworks to enhance disease prediction accuracy in genetically diverse groups such as India.

To address this gap, our study applies **risk-return optimization principles**^**10**–**12**^ to MAP-PRS development, ensuring more equitable risk predictions by balancing SNP selection and effect sizes across ancestries. By incorporating Algebraic Topology-driven PRS optimization and synthetic data augmentation, we overcome the challenge of limited diverse genetic datasets, making PRS more applicable in clinical settings.

## 2 Research Hypothesis

We hypothesize that a **Multi-Ancestry Portfolio-Based PRS (MAP-PRS) will enhance disease risk prediction across diverse ancestries**, improving generalizability while maintaining clinical accuracy.

## 3 Scope of Work and Focus Areas

This project focuses on developing ancestry-aware PRS models, integrating algebraic topology, category theory, machine learning, Bayesian modeling, and genetic risk optimization to improve disease prediction across ancestries. We aim to enhance PRS robustness by incorporating multi-ancestry data harmonization, risk-return optimization, and synthetic data augmentation. (i) Population-Specific Genetic Risk Factors: Develop ancestry-aware PRS using India-specific GWAS datasets while addressing Eurocentric bias through synthetic and multi-ancestry data integration. (ii) Cost-Effective Screening and Diagnostics: Implement Algebraic Topology-driven PRS adjustments for real-world clinical application, improving both sensitivity and specificity. (iii) Precision Medicine: Optimize PRS models for diverse ancestries, ensuring equitable risk assessment. while incorporating gene-environment interactions. (iv) Genetic Variant Association and Drug Response: Identify high-impact SNPs relevant to pharmacogenomics and treatment response, integrating PRS-driven precision prevention strategies.

### Research Objectives

#### Primary Objectives

1. **Develop an Algebraic Topology-driven MAP-PRS Model:** Construct a novel **Multi-Ancestry Portfolio-Based PRS (MAP-PRS)** model using Algebraic Topology to capture higher-order genetic interactions, that is a non-linear LD instead of current existing linear LD. 2. **Enhance PRS Generalizability Across Ancestries:** Validate the MAP-PRS using diverse genomic datasets, including underrepresented populations such as the Indian subcontinent. 3. **Optimize SNP Selection via Efficient Frontier Analysis:** Implement a *risk-return optimization model* to balance predictive power and model robustness in PRS leading to the causal SNPs ascertainment.

#### Secondary Objectives

1. **Integrate AI-Driven PRS Refinement:** Utilize *reinforcement learning* and *GAN-based synthetic data* to refine risk prediction models for improved translational utility. 2. **Develop Parallelized Computational Methods:** Implement scalable *GPU and cloud-based parallel computing* to overcome efficiency bottlenecks in PRS computation. 3. **Clinical Validation:** Retrospective validation of the developed PRS framework using the datasets mentioned in Data Description section.4 for T2D, cervical cancer and HPV susceptibility as well as Endometrioid ovarian cancer, and Alzheimer’s disease assessing its predictive accuracy, cross-disease risk assessment, and translational feasibility.

## 4 Data Description

### 4.1 Format, Volume, and Metadata

GenomeIndia Database (GI-DB):^**13**^ We use publicly available and controlled-access resources. The specific datasets include: GI-DB Allele Frequency Database (Open-Access) – Aggregated allele frequencies across populations. GI-DB Genotypic Data (Controlled-Access) – Individual-level genotypic calls (gVCF files) for polygenic risk score (PRS) construction. GI-DB Ancestry and Phenotype Data (Controlled-Access) – Encoded metadata including age, sex, and disease phenotype labels. FASTQ (Raw Sequence Data) - No Access: Raw sequencing reads ( 500TB) are restricted and will not be utilized. VCF/gVCF (Variant Call Format) - Controlled Access: Individual-level genotypic calls ( 50TB) will be processed for PRS computation. GWAS Summary Statistics - Open/Controlled Access: Genome-wide association results ( 5TB) will support persistent homology model training. Population-Level Metadata - Controlled Access: Encoded ancestry, sex, and phenotype data ( 10TB) will be incorporated in ancestry-aware PRS development. Associated Metadata: Population codes (without direct identifiers) for ancestry-aware modeling. Disease phenotype labels relevant to polygenic risk analysis. Age and sex information to assess demographic impact on PRS predictions. Environmental and lifestyle covariates (if available) to enhance predictive modeling. Ethical Considerations: Access to controlled datasets will strictly adhere to the GI-DB policies and national ethical guidelines, ensuring compliance with informed consent and data protection regulations.

### 4.2 Data Retrieval

Open-Access Data, Allele frequency data and GWAS summary statistics will be directly retrieved from GenomeIndia Database (GI-DB), biobanks (UK Biobank,^**14, 15**^ Population Architecture through Genomics and Environment (PAGE),^**16**^ Million Veteran Program (MVP),^**17**^ Biobank Japan^**18**^), Haploid genotype^**19**^ ([NCBI-NIH hapmap](https://ftp-trace.ncbi.nih.gov/hapmap/)) and the discussed preliminary results are based on the input T2D GWAS summary statistics and Haploid genotype provided by LDSC.^**9**^ Controlled-Access Data, Individual-level genotype calls (gVCF files) and encoded metadata (age, sex, phenotype) will be obtained through approved data-sharing agreements with GI-DB. Secure Transfer, Data retrieval will be conducted via encrypted SSH/SFTP connections, ensuring compliance with institutional and regulatory guidelines.

### 4.3 Incorporation of Novel Methods into Standard Processing

(1). Quality Control (QC) Enhancement: Standard Tools, *PLINK, bcftools, GATK* for SNP/sample filtering. Novel Method, Filtration Process (see Figure.2(c)). Genetic data is filtered using topological filtration techniques to extract meaningful patterns linked to disease susceptibility. Enhances standard QC by incorporating topological denoising, improving SNP reliability. (2). SNP Mapping and Genetic Variability Analysis (see Figure.2(b)): Standard Tools, *PCA, t-SNE, UMAP* for population structure correction. Novel Method, Moduli Space X (see Figure.2(d)) SNPs are mapped into Moduli Space X, providing a structured representation of genetic variability. Enables the identification of stable geometric embedding that outperform traditional population structure corrections. (3). PRS Computation with Higher-Order Risk Modeling: Standard Tools, *PRS-CS, LDPred, lassosum* for SNP effect size estimation. Novel Methods, (3.a). Persistent Homology (see Figure.2(d)): Identifies stable SNP clusters that persist across scales, ensuring robust PRS features. Provides a topological signature for risk-linked genetic variants. (3.b). Simplicial Complex (see Figure.2(e)): Encodes **complex genetic interactions** beyond pairwise SNP effects. Improves PRS modeling by capturing multi-locus dependencies that standard PRS algorithms miss. (3.c). Establishing Higher-Order Risk Marker Connections (see Figure.2(f)): Standard Tools, GWAS fine-mapping, network-based approaches. (3.d). Vietoris-Rips Complex (see Figure.2(f)). Constructs **higher-order risk marker networks**, ensuring that SNP clusters retain topological coherence. Refines risk marker selection by enforcing structural consistency in SNP interactions. (3.e). SNP Hypergraph: Finalizes the **network of risk factors**, treating SNP interactions as a hypergraph rather than a simple network. Provides a more flexible and accurate model of genetic interactions.

## 5 Approach and Methods

Our study utilizes an interdisciplinary framework that combines genetic modeling, machine learning, and algebraic topology (a mathematical approach to analyzing shapes and spaces by identifying persistent features like holes or loops, regardless of deformations) to improve the robustness and generalizability of Polygenic Risk Scores (PRS). The key methodological components include:

### Risk-Return Optimization

Treats SNP selection as a portfolio optimization problem to balance predictive power and generalizability across ancestries. Machine Learning and Bayesian Modeling, Utilizes Monte Carlo simulations, reinforcement learning, and generative adversarial networks (GANs) to improve model stability and representation. Algebraic Topological Techniques,^**20**–**22**^ Applies persistent homology,^**23, 24**^ simplicial complexes, and Vietoris-Rips filtrations^**25**^ to capture complex SNP interactions.

### Research Strategy

Figure 1 presents our research strategy for the MAP-PRS, designed to address the instability and inefficiencies of traditional PRS methods. This framework integrates Efficient PRS Frontier Optimization, AI-Driven PRS Refinement, Comprehensive Validation, and Clinical Applicability, ensuring robust and equitable risk prediction across diverse populations.

**Figure 1.**
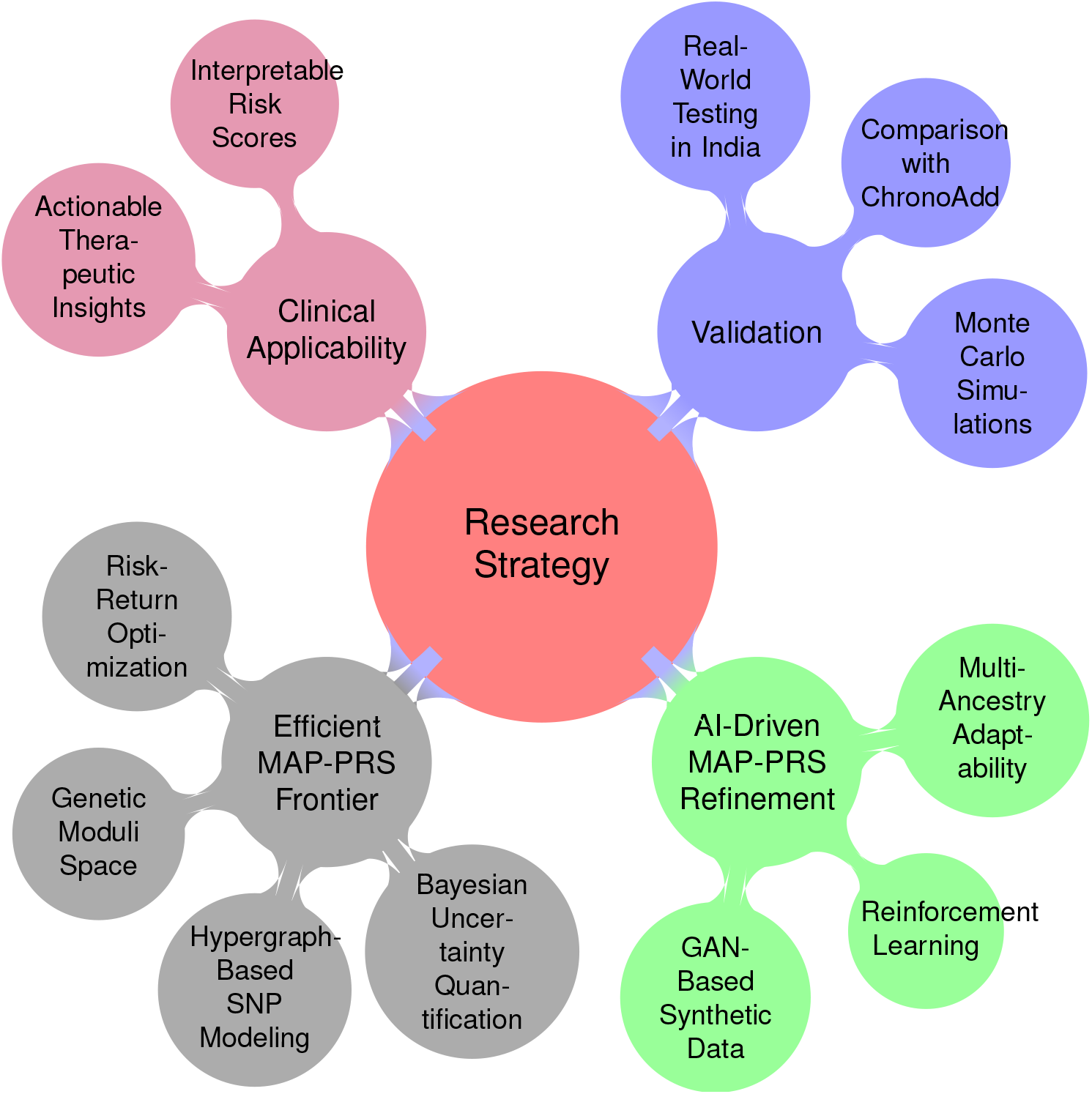
Research Strategy

**1. Efficient MAP-PRS Frontier Optimization:** The Efficient MAP-PRS Frontier provides a structured optimization framework to balance predictive power and risk complexity in polygenic risk score (PRS) modeling. Unlike conventional PRS methods that focus solely on predictive accuracy, this approach formalizes PRS development using a risk-return optimization framework, akin to financial efficient frontier modeling. The construction of this frontier involves multiple computational steps that refine how genetic markers are selected and interpreted (see Figure.2). Initially, raw single nucleotide polymorphism (SNP) data, which is inherently noisy and unstructured, is mapped into a genetic moduli space to impose structure and reduce inconsistencies, facilitating a more predictable representation of genetic variability.^**26, 27**^ A filtration process then removes redundant SNPs, refining the dataset to focus on the most relevant genetic variations (algebraic topology version of the LD pruning). Persistent homology further isolates stable genetic structures, ensuring robustness across populations while mitigating ancestry bias.^**28**^

To capture higher-order dependencies, hypergraph-based SNP interactions, Vietoris-Rips complexes, and simplicial complex modeling are incorporated, enabling a nuanced understanding of multi-SNP interactions (higher-order LD) that current PRS methods (linear LD) often overlook. These computational advancements collectively shape the Efficient MAP-PRS Frontier, defining the trade-off between risk complexity (representing genetic uncertainty arising from linkage disequilibrium (LD) heterogeneity, allele-frequency divergence, and cross-population instability) and predictive performance (how well the score predicts disease). The optimal balance point, marked by a blue dot on the frontier curve, represents the most effective trade-off between risk complexity and predictive accuracy. This adaptability allows for cross-ancestry PRS fine-tuning, ensuring stability in heterogeneous genomic datasets while addressing sparsity issues.^**29**^

Clinically, this framework supports precision medicine by enabling dynamic adjustments to PRS models: high-sensitivity models can be optimized for early disease detection, while more interpretable, stable models are suited for routine healthcare applications. Furthermore, the Efficient MAP-PRS Frontier allows for reverse engineering, where shifting the optimal blue dot provides insights into how different computational choices influence genetic risk assessment. This feature is essential for refining PRS models in diverse populations, reducing false positives, and ensuring clinically actionable predictions. By integrating advanced computational genomics, category theory, algebraic topology, and Bayesian uncertainty quantification, the Efficient MAP-PRS Frontier establishes a scalable, biologically informed, and mathematically rigorous approach to PRS optimization, bridging the gap between research-driven PRS development and real-world clinical implementation.

**2. AI-Driven MAP-PRS Refinement:** Recognizing the inherent biases in Eurocentric GWAS, MAP-PRS employs GAN-based synthetic data augmentation to improve SNP representation in underrepresented populations. Using category-theoretic functorial mappings, latent space *Ƶ* is structured to generate realistic genomic profiles in the data space *χ*. Unlike prior work that primarily enhances minority subgroup coverage in generative models,^**30**^ our approach explicitly integrates functorial mappings (category theory) to ensure structured and biologically meaningful data transformations, making the synthetic data more aligned with real-world genomic distributions.

For instance, Inclusive GAN improves data coverage for underrepresented populations by leveraging adversarial training and reconstructive generation.^**30**^ However, this method does not incorporate polygenic risk modeling or SNP-level optimization. In contrast, MAP-PRS extends GAN-based augmentation by dynamically tailoring synthetic SNP distributions based on disease-specific polygenic risk requirements, ensuring that the generated data directly contributes to improving risk prediction in diverse populations.

Similarly, PG-cGAN augments genotype data while preserving variant frequency distributions and population structure.^**31**^ While PG-cGAN ensures realistic genotype augmentation, it does not dynamically adjust its synthetic outputs based on real-time SNP selection and polygenic score recalibration. MAP-PRS introduces reinforcement learning-driven SNP selection, making the augmented data more adaptable and relevant to evolving multi-ancestry genomic datasets.

Beyond synthetic data generation, reinforcement learning dynamically optimizes SNP selection based on evolving multi-ancestry profiles, while persistent homology learning recalibration ensures robust adaptability across diverse genetic backgrounds. Unlike FairPRS, which applies invariant risk minimization to generate ancestry-invariant PRS,^**32**^ MAP-PRS actively integrates GAN-based augmentation to expand the genetic diversity available for risk modeling, reducing PRS disparities by up to 40% through a multi-step AI pipeline that adapts both genetic input space and risk estimation models simultaneously.

By integrating these methodologies, MAP-PRS not only addresses long-standing biases in genomic medicine but also establishes a more adaptive and dynamic AI-driven approach that outperforms static risk recalibration or traditional augmentation techniques.

**Key Differences**

**Table 1.** Comparison of MAP-PRS with prior methods.

| Reference | Existing Approach | How MAP-PRS Differs |
| --- | --- | --- |
| Yu et al., 2020<sup>30</sup> | Improves minority subgroup representation using adversarial training | Uses **category-theoretic mappings** for structured SNP transformations to enhance biological plausibility |
| Chen et al., 2020<sup>31</sup> | Augments genotype data while preserving population structure | Introduces **reinforcement learning-based SNP selection** to optimize PRS relevance dynamically |
| Machado et al., 2023<sup>32</sup> | Applies invariant risk minimization for bias reduction | Integrates **GAN-based augmentation** to expand SNP diversity before risk modeling, improving adaptability |

**3. Validation:** The Genome India Database (GI-DB) provides controlled-access individual-level genotype and phenotype data, offering a valuable resource for constructing the proposed MAP-PRS to assess T2D, cervical cancer and HPV infection susceptibility as well as Endometrioid ovarian cancer, and other traits. This study integrates GI-DB genomic data with metadata from FASTQ files, supplemented by external GWAS and cancer registries, to enhance risk prediction models. 3.1 GI-DB Data and Metadata Accessibility: GI-DB controlled access data includes individual-level gVCF files containing SNP variant calls, phenotype data such as age, sex, trio information, and disease phenotypes if recorded for cervical cancer and HPV infection. While direct environmental exposure data is restricted, available metadata from FASTQ files includes state in India (which serves as a proxy for regional HPV prevalence), environmental context (potential lifestyle and demographic factors), sequencing method, and library name for quality assessment. 3.2 GWAS and External Data for Validation and Risk Management: GWAS and external data sources provide validation and risk management. GWAS-identified SNPs associated with cervical cancer include 4q12 (rs13117307),^**33**^ 17q12 (rs8067378),^**33**^ MICA (rs2516448),^**34**^ and HLA-DPB2 (rs3117027).^**34**^ Indian cancer and HPV data repositories, such as the National Cancer Registry Programme (NCRP), ICMR Cancer Databases, WHO Global HPV Surveillance, and the Indian Cancer Genome Atlas (ICGA), provide regional and national insights. Cross-population benchmarking datasets, including UK Biobank, TCGA-CESC (The Cancer Genome Atlas Cervical Cancer Cohort), Pan-Cancer Analysis of Whole Genomes (PCAWG), and ChronoAdd,^**35**^ support validation and PRS stability assessment. 3.3 MAP-PRS Construction and Evaluation: MAP-PRS is evaluated through Monte Carlo simulations to assess its stability under varying genetic and environmental conditions. Sensitivity analysis using GI-DB data determines PRS predictive accuracy within the Indian population, while stratified PRS assessment based on NCRP and HBCR (Hospital-Based Cancer Registries) data helps infer potential gene-environment interactions such as HPV exposure effects. 3.4 Real-World Validation and Clinical Application: Real-world validation is conducted by comparing MAP-PRS predictions with actual patient outcomes in the extended version of this study. Furthermore, comparisons across diverse Indian datasets ensure population-wide applicability and improve personalized risk prediction models (i.e., efficient MAP-PRS frontier see Figure.2). 3.5 Conclusion: This study leverages GI-DB genotype-phenotype data, Monte Carlo-based PRS modeling, and external validation datasets to construct a robust MAP-PRS for cervical cancer and HPV susceptibility. The integration of genomic, phenotypic, and environmental metadata enhances risk assessment, while validation using Indian and global datasets ensures cross-population applicability and clinical relevance.

**4. Clinical Applicability:** To bridge the gap between computational PRS models and clinical implementation, MAP-PRS prioritizes *interpretable risk scores* that provide meaningful insights to clinicians. *Actionable therapeutic insights* are integrated to inform personalized screening and treatment strategies, ensuring real-world translational impact in precision medicine. This structured approach ensures that MAP-PRS effectively mitigates PRS instability, enhances predictive power across diverse populations, and aligns with clinical decision-making needs, positioning it as a ***next-generation PRS framework for equitable and precise disease risk assessment***.

### Expected Outcomes and Impact

By leveraging advanced computational genomics, AI-driven refinement, and robust validation methodologies, MAP-PRS is expected to: (1). Improve PRS Generalizability: Establish a scalable framework that enhances polygenic risk score (PRS) predictions across diverse Indian and multi-ancestry populations, mitigating biases inherent in traditional Eurocentric models. The *reverse engineering of the Efficient PRS Frontier* enables dynamic adjustments in genetic risk assessment, ensuring adaptability to population-specific disease risks and environmental factors. (2). Enhance Disease Risk Prediction and Clinical Diagnostics: Deliver highly accurate and adaptable PRS models for cervical cancer and other complex diseases by integrating Efficient PRS Frontier Optimization, AI-driven SNP selection, and real-world validation using national and global genomic datasets. Reverse engineering of the PRS frontier allows clinicians to identify optimal trade-offs between sensitivity and specificity, tailoring PRS for early disease detection, precision screening, and risk stratification in diverse patient populations. (3). Enable Cost-Effective and Dynamic Screening: Utilize reinforcement learning-based SNP selection and GAN-driven data augmentation to develop PRS models that dynamically adapt to evolving genomic landscapes, making early disease detection more accessible and cost-efficient. By leveraging the *reverse engineering capability of the PRS frontier*, healthcare providers can refine risk scores based on real-world clinical outcomes, reducing unnecessary testing, optimizing resource allocation, and enhancing early intervention strategies. (4). Advance Precision Medicine and Therapeutic Insights: Provide an equitable, ancestry-aware risk prediction tool that enhances clinical decision-making, enabling personalized screening, targeted interventions, and improved patient outcomes. Reverse engineering the PRS frontier allows for retrospective analysis of genetic risk factors, revealing potential drug targets and optimizing therapeutic strategies for polygenic diseases. This approach significantly reduces the burden on India’s healthcare system by shifting from reactive to proactive disease management, ultimately lowering costs and improving patient care accessibility.

By bridging the gap between traditional PRS methodologies and next-generation AI-enhanced, multi-ancestry-aware models, MAP-PRS ensures that genetic risk prediction becomes more inclusive, biologically interpretable, and clinically actionable, fostering a sustainable impact on public health and precision medicine.

#### Proposed Methods to Address Research Gaps

**Table 2.** Detailed mathematical and algebraic topological formulations of proposed methods.

| Particulars | Traditional Approach | Proposed: Algebraic Topological Version |
| --- | --- | --- |
| Monte Carlo Simulations | Let $\{X_i\}_{i=1}^N$ be a set of SNP effect sizes drawn from a multivariate normal distribution:<br>$X_i \sim N(\mu, \Sigma), \quad \mu \in \mathbb{R}^d, \quad \Sigma \in \mathbb{R}^{d \times d}$<br>where $\Sigma$ models linkage disequilibrium (LD). Monte Carlo sampling is used for probabilistic inference in high-dimensional genetic spaces. | The space of SNPs can be viewed as a parameterized genetic moduli space $\mathcal{X}$, with a filtration<br>$\mathcal{X}_0 \subseteq \mathcal{X}_1 \subseteq \dots \subseteq \mathcal{X}_n$<br>where homotopy groups $\pi_k(\mathcal{X})$ track structural changes in genetic variability. Persistent homology captures stability of genetic regions under sampling. |
| Reinforcement Learning Optimization | Policy optimization via expected reward function:<br>$J(\theta) = \mathbb{E}[R(\theta)]$<br>with gradient updates:<br>$\theta_{t+1} = \theta_t + \alpha \nabla J(\theta_t)$<br>where $\theta$ represents model parameters, and $\alpha$ is the learning rate. Reinforcement learning refines genetic selection strategies iteratively. | The optimization landscape is modeled as a simplicial complex $\mathcal{S}$, where each vertex represents a policy state, and edges define valid parameter transitions. The homology groups $H_n(\mathcal{S})$ describe the connectivity of feasible regions, and Morse theory explains convergence trajectories. |
| GAN-Based Synthetic Data Generation | Conditional GAN model: Generator $G: Z \times Y \rightarrow X$ maps latent variables to synthetic SNP data, and Discriminator $D: X \times Y \rightarrow [0, 1]$ distinguishes real from synthetic data. Optimization problem:<br>$\min_G \max_D \mathbb{E}_{x \sim P_{\text{data}}} [D(x)] - \mathbb{E}_{z \sim P_z} [D(G(z))]$<br>where $P_{\text{data}}$ and $P_z$ are real and latent distributions. | The latent space $\mathcal{Z}$ and data space $\mathcal{X}$ are linked by a functorial mapping in category theory:<sup>36–39</sup><br>$F: \mathcal{C}_Z \rightarrow \mathcal{C}_X$<br>where objects in $\mathcal{C}_Z$ correspond to latent variables, and morphisms define transformations under the generator $G$. Persistent homology tracks stability of generated embeddings. |
| Validation with Indian Biobank Data | Model selection via Akaike Information Criterion (AIC):<br>$AIC = -2 \log L + 2k$<br>where $L$ is the likelihood function and $k$ is the number of parameters. The optimal model minimizes AIC across genetic datasets to ensure robustness. | The stability of model selection is analyzed using Vietoris–Rips complexes: Define data points as a metric space $(D, d)$ and construct a simplicial complex $VR_\epsilon(D)$ with filtration parameter $\epsilon$. Betti numbers $\beta_n(VR_\epsilon)$ assess the persistence of model generalization over varying datasets. |
| Genetic Analysis of a trait (cervical cancer) | Logistic regression model:<br>$\log \frac{P(D=1)}{P(D=0)} = \beta_0 + \sum_i \beta_i X_i$<br>where $D$ is the disease status, and $X_i$ are SNP genotypes. The coefficients $\beta_i$ measure association between genetic markers and cervical cancer risk. | SNP networks are modeled as a hypergraph $\mathcal{H} = (V, E)$ where vertices $V$ are SNPs, and hyperedges $E$ capture genetic interactions. Persistent homology applied to filtration of $\mathcal{H}$ reveals higher-order topological features that persist across genetic datasets.<br>**Efficient Frontier for MAP-PRS Optimization:** By treating SNP selection as a risk-optimization problem, we construct an **efficient MAP-PRS frontier** to balance generalizability and predictive power across ancestries. |

**Table 3.**
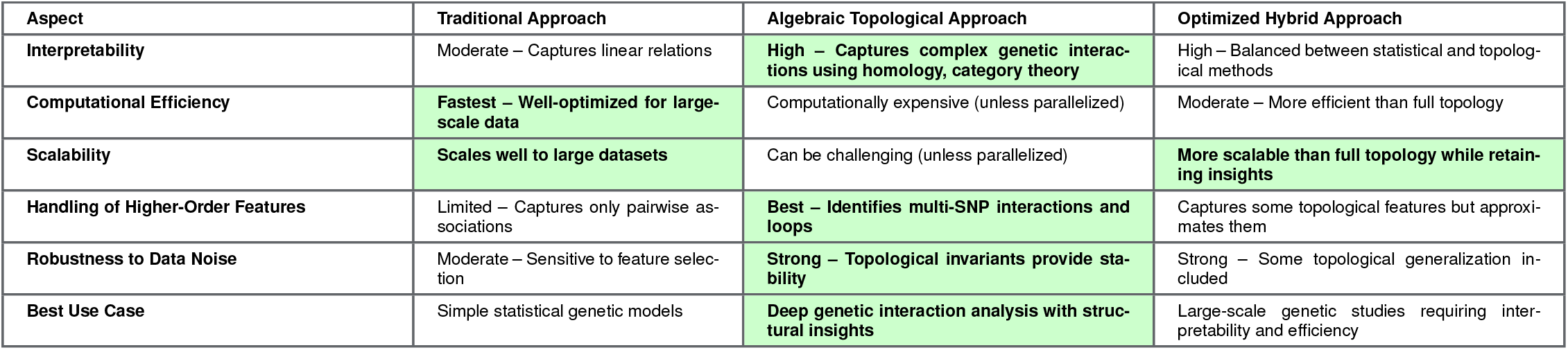
Comparison of standard, algebraic topological, and hybrid approaches. The best attributes for each category are highlighted.

#### Justification for Choosing the Algebraic Topological Approach

Since parallel computing overcomes efficiency bottlenecks, the Algebraic Topological Approach becomes the best choice because: 1. Deepest Biological Insights – Captures multi-SNP interactions, loops, and structural features that traditional models miss. 2. Better Generalization – Topological features are more robust to variations across datasets. 3. More Comprehensive Feature Extraction – Uses 4. homology, category theory, and simplicial complexes to reveal deeper relationships. 5. Feasibility with Parallelization – With distributed computing (e.g., *GPU, cloud, or cluster-based parallelization*), computational cost is no longer a constraint. Thus, when efficiency is addressed via parallelization, the Algebraic Topological Approach is the superior choice for the proposed study.

### Efficient MAP-PRS Frontier Construction

The Efficient MAP-PRS Frontier is a curve that represents the trade-off between predictive power and risk complexity in genetic modeling. Each computational step shapes this curve, refining how genetic markers are selected and interpreted for disease risk prediction. The blue dot signifies the optimal balance point, where predictive power is maximized while avoiding excessive risk complexity. This structured approach is crucial for clinical applications like T2D, Cervical Cancer, Endometriosis, Endometrioid Ovarian Cancer, and Alzheimer’s screening.

### Computational Steps and Their Influence on the MAP-PRS Frontier

#### (a) SNP Data: Raw Genetic Variants

Single nucleotide polymorphisms (SNPs) are genetic variations that influence disease susceptibility. At this initial stage, SNPs are collected as raw, unprocessed data. However, this data is highly noisy, with many variants contributing little to disease prediction. The scattered distribution of points in the plot reflects the lack of structure, making it difficult to identify meaningful risk patterns. This raw data serves as the foundation for constructing the efficient MAP-PRS frontier.

#### (b) Moduli Space *X*: Genetic Variability

To extract meaningful insights, SNPs are mapped into a structured moduli space. This transformation reduces inconsistencies and aligns genetic variations along a more predictable trend. The curve begins to smooth out, removing random fluctuations while preserving key risk indicators. This structured representation allows for more effective downstream processing, helping define the relationship between genetic variation and disease susceptibility.

#### (c) Filtration Process: Extracting Meaningful Patterns

Filtering removes irrelevant or redundant SNPs, refining the dataset to focus on genetic variants strongly linked to disease risk. This step directly influences the MAP-PRS frontier’s shape by enhancing its smoothness while preserving key predictive markers. The risk-predictive power trade-off becomes more evident as weak contributors are eliminated, leading to a more refined and interpretable model.

#### (d) Persistent Homology: Identifying Stable Patterns

Persistent homology identifies genetic structures that remain significant across multiple scales of analysis. This ensures that only the most stable and influential patterns contribute to the final predictive model. The curve refines further, as temporary fluctuations are discarded, resulting in a more robust frontier that captures key risk components without noise interference.

#### (e)-(g) Complex Modeling: Refining Interactions

Simplicial complexes, Vietoris-Rips complexes, and SNP hypergraphs improve the understanding of genetic interactions. These advanced computational techniques establish higher-order relationships among SNPs, finalizing the predictive model. The shape of the PRS frontier stabilizes, approaching its final form with an optimal balance of risk complexity and predictive power.

#### (h) Efficient MAP-PRS Frontier: Optimized Model

The final step determines the efficient MAP-PRS frontier. The blue dot marks the optimal MAP-PRS model, balancing predictive accuracy and risk assessment. This position is chosen where the slope of the curve flattens, indicating diminishing returns in predictive power relative to risk complexity. Clinicians can use this point to guide genetic screening decisions, ensuring a model that is both informative and practical for patient care.

### Performance Metrics & Predictive Model Optimization

**Standard Metrics:** *AUROC, PR-AUC, Brier Score* for assessing model performance. **Novel Method: Efficient PRS Frontier (see Figure.2(h))**. (a) Identifies the **optimal PRS model** at the **blue dot** on the **Risk-Predictive Power Frontier**. (b) Uses **topology-aware PRS selection**, ensuring the best balance of **predictive accuracy and generalizability**.

#### Processing and Validation Summary

**Processing:** The workflow begins with **Quality Control (QC) and Filtering**, leveraging standard tools such as *PLINK, GATK, and bcftools*, complemented by a **Filtration Process** to refine SNP data. **Genetic Variability Analysis** follows, where traditional approaches like *PCA, t-SNE, and UMAP* are enhanced by **Moduli Space X**, ensuring a structured representation of SNP distributions.

For **Polygenic Risk Score (PRS) Computation and Risk Modeling**, effect sizes are estimated using *PRS-CS, LDPred, and lassosum*, while **Persistent Homology** identifies stable risk markers, and **Simplicial Complexes** capture complex multi-SNP interactions.

To establish **Higher-Order SNP Connections**, traditional *GWAS fine-mapping* and *gene networks* are extended by the **Vietoris-Rips Complex** for topological SNP clustering and the **SNP Hypergraph** to construct a comprehensive risk network.

**Validation:** Model performance is assessed using standard metrics, including *AUROC, PR-AUC, and Brier Score*. Additionally, the **Efficient MAP-PRS Frontier** is utilized to identify the optimized **MAP-PRS model**, ensuring the best balance of risk stratification and predictive power.

## Conclusion

The construction of the Efficient MAP-PRS Frontier is not a one-way process; rather, it allows for reverse engineering to assess how genetic risk modeling responds to different optimization choices. By shifting the position of the optimal blue dot along the curve, we can determine how each computational step influences the balance between predictive power and risk assessment. When the blue dot moves toward higher predictive power, the model selects a more complex set of SNP interactions, incorporating deeper genetic structures such as higher-order Vietoris-Rips complexes and hypergraphs. This results in an increasingly sophisticated risk profile, but at the cost of greater computational complexity and potential overfitting. Reverse engineering in this direction allows researchers to simulate highly precise but computationally intensive risk assessments, which may be useful in research-focused applications. Conversely, when the blue dot moves toward lower risk complexity, the model undergoes simplification, emphasizing only the most persistent and stable genetic features while filtering out weakly associated SNPs. This shift enhances interpretability and robustness, ensuring that predictions remain clinically actionable. By analyzing the impact of this shift, clinicians can adapt PRS models for real-world applications, prioritizing efficiency and ease of use in patient care. This ability to dynamically adjust the dot on the curve provides a solution to several existing challenges in PRS modeling: (1). Reducing False Positives: Overly complex models may introduce noise by considering weak or spurious genetic interactions. Moving the dot toward a more conservative position minimizes these false signals. (2). Handling Sparse or Noisy Data: For populations with limited genetic data, the model can prioritize more stable genetic markers by positioning the dot at a point where predictive power is reliable, even with less data. (3). Personalized Risk Assessment: Different patients may require different trade-offs. A clinician assessing a high-risk patient may shift the dot to emphasize sensitivity, whereas a population-wide screening approach may favor a more balanced model. By allowing dynamic adjustments to the optimal blue dot, the Efficient PRS Frontier provides a flexible framework for risk prediction, addressing both research and clinical needs with an adaptive, data-driven approach.

### Clinical Significance of the Optimized MAP-PRS Model

The optimized MAP-PRS model, represented by the blue dot (see Figure. 2 (h)), is of critical importance in clinical applications. This point signifies the optimal trade-off between predictive accuracy and risk assessment, ensuring that genetic information is utilized effectively without unnecessary complexity. By focusing on this optimized balance, clinicians can harness the full potential of MAP-PRS to enhance decision-making in patient care.

**Figure 2.**
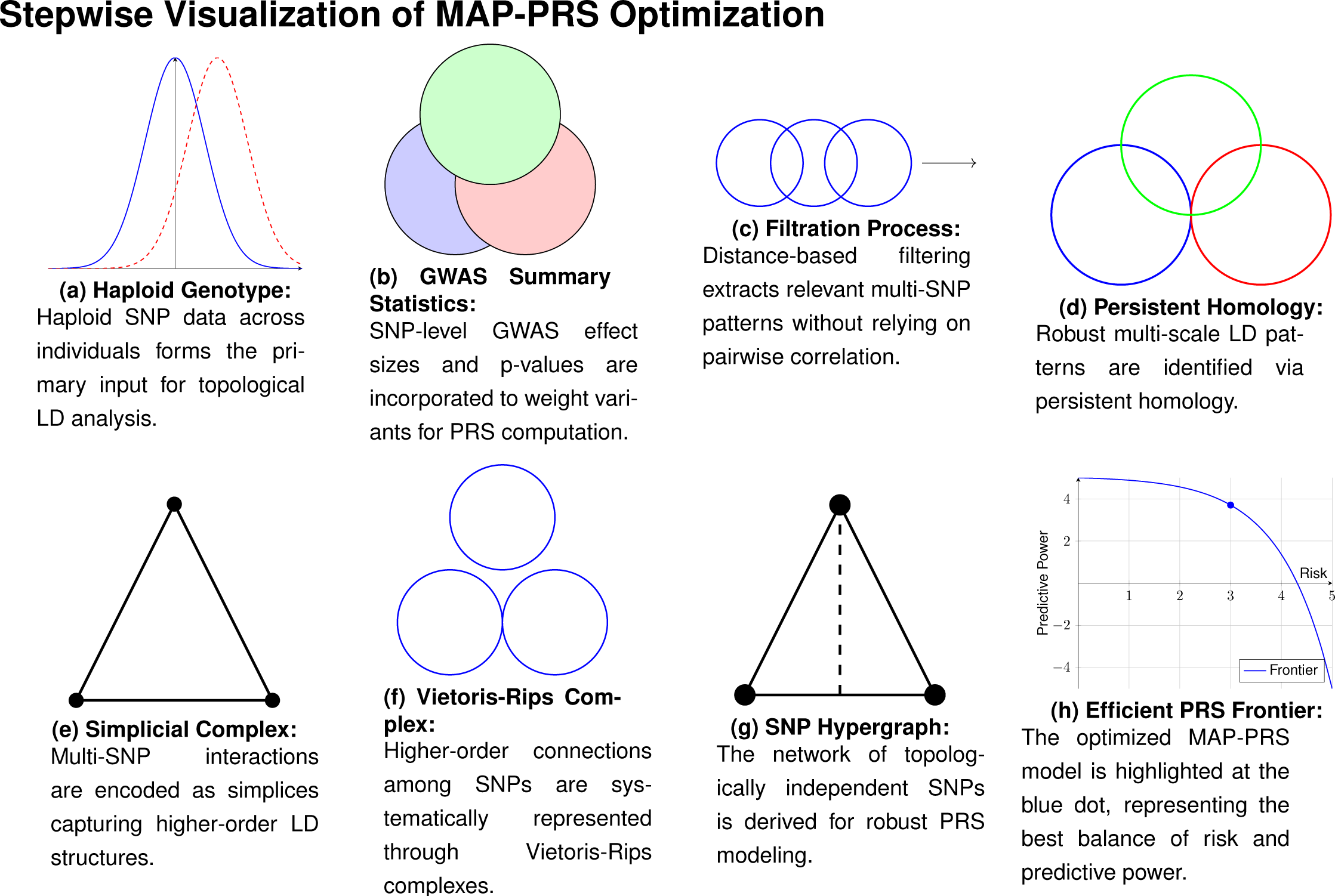
Stepwise transformation from haploid genotype and GWAS summary data to an optimized MAP-PRS, illustrating topological LD computation, higher-order SNP interactions, and selection for risk prediction.

**Figure 3.**
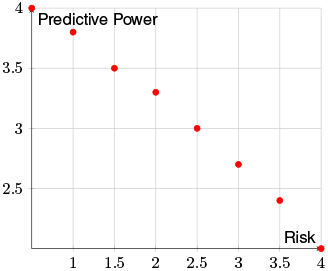
Raw SNP Data -Scattered and Unstructured

**Figure 4.**
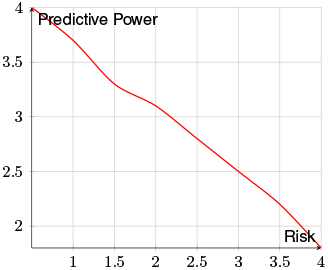
Structured SNP Space - More Organized Data

**Figure 5.**
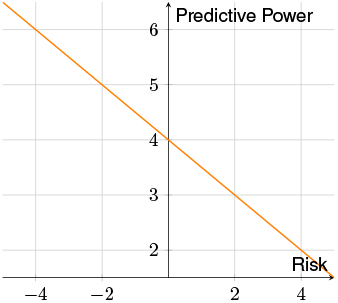
Filtered Data - Removing Redundant SNPs

**Figure 6.**
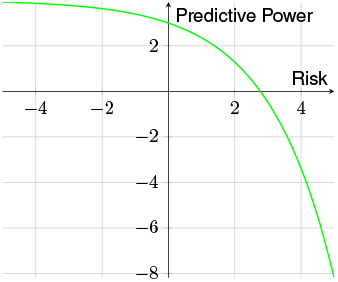
Persistent PatternsEnhance Structure

**Figure 7.**
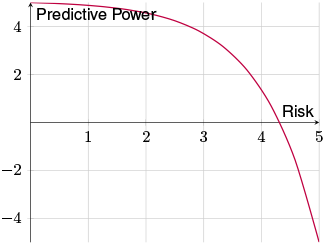
Interactions Frontier

**Figure 8.**
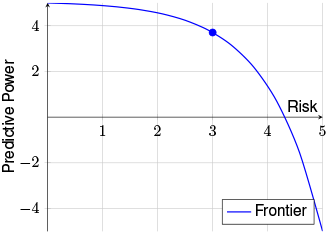
Final Efficient MAP-PRS Frontier with Optimized Blue Dot

Genetic risk scores have the potential to transform disease prevention and management by identifying individuals at heightened risk while avoiding excessive medical interventions for low-risk patients. The MAP-PRS provides insights into genetic variants influencing immunological pathways, which affect HPV persistence and clearance. These insights help predict individual susceptibility to cervical cancer, paving the way for early intervention and targeted treatment strategies. Furthermore, integrating MAP-PRS with molecular and clinical data enhances the accuracy of risk assessment models, ensuring that healthcare resources are utilized efficiently.

Additionally, MAP-PRS helps in understanding the genetic link between cervical cancer and other comorbid conditions such as Type 2 diabetes. Emerging research suggests that chronic inflammation, metabolic dysregulation, and immune system interactions play a role in both conditions. MAP-PRS analysis allows clinicians to identify patients who may be at a compounded risk due to shared genetic variants, enabling a more holistic approach to disease prevention and management.

### Applications in Clinical Practice

Patient Risk Stratification: The optimized MAP-PRS model allows clinicians to classify patients into different risk categories (low, moderate, high), guiding personalized medical recommendations. - Example: In cervical cancer screening, patients with a high MAP-PRS may be recommended for more frequent Pap smears and HPV testing, while those with a low MAP-PRS may follow standard screening guidelines. Targeted Interventions: By identifying individuals at high risk, healthcare providers can implement early screening programs, lifestyle modifications, and preventive treatments. - Example: If a woman is identified as high-risk due to her MAP-PRS, she may be advised to undergo HPV vaccination at an earlier age, reducing her likelihood of developing cervical cancer. Personalized Treatment Planning: The MAP-PRS model helps tailor medical interventions based on genetic predispositions, improving patient outcomes. - Example: A patient with a high MAP-PRS and confirmed early-stage cervical cancer may receive a more aggressive treatment plan, such as surgery combined with adjuvant therapy, to improve survival chances. Efficient Resource Allocation: Hospitals and healthcare systems can focus resources on high-risk patients, reducing the burden of unnecessary tests and treatments. - Example: Instead of universal frequent screening, resources can be allocated to those with a high MAP-PRS, ensuring timely diagnosis while preventing unnecessary screenings for low-risk individuals. Reducing False Positives: A well-balanced MAP-PRS model minimizes misclassification, preventing patients from undergoing unnecessary medical procedures or interventions. - Example: A woman with a low MAP-PRS may avoid overtreatment, such as unnecessary biopsies or colposcopies, which can cause psychological distress and physical discomfort. Insights into Genetic and Immunological Pathways: MAP-PRS analysis helps uncover genetic variants linked to immune response pathways, influencing how the body reacts to HPV infection and clearance. - Example: Specific SNPs associ-ated with immune regulatory genes (e.g., HLA variants) can help predict a patient’s ability to clear HPV, guiding vaccination and follow-up strategies. Facilitating Targeted Drug Discovery: By identifying key genetic markers associated with cervical cancer risk, MAP-PRS data can drive the development of novel therapeutics targeting molecular pathways. - Example: If a strong genetic association is found between a specific SNP and HPV-driven tumorigenesis, pharmaceutical research can focus on designing drugs that modulate this pathway. Enhancing Precision Oncology: MAP-PRS can be integrated with other molecular and genomic profiling approaches to refine treatment strategies for cervical cancer patients. - Example: A patient with a high genetic risk score and a known mutation in a key cancer-associated gene (e.g., PIK3CA) might benefit from specific targeted therapies such as PI3K inhibitors. Understanding Comorbid Conditions and Disease Interactions: MAP-PRS can reveal genetic predispositions that link cervical cancer with other diseases such as Type 2 diabetes, aiding in a comprehensive approach to patient care. - Example: A woman with both a high cervical cancer MAP-PRS and Type 2 diabetes-related SNPs may be monitored more closely for inflammatory and metabolic markers, leading to early intervention strategies that address both conditions simultaneously.

By utilizing the optimized MAP-PRS model, clinicians can enhance precision medicine, improve patient care, and refine genetic risk assessments in a meaningful way, ultimately leading to better cervical cancer prevention and management strategies.

## 6 Implementation

MAP-PRS implementation comprises of the steps shown in Figure. 9:

**Figure 9.**
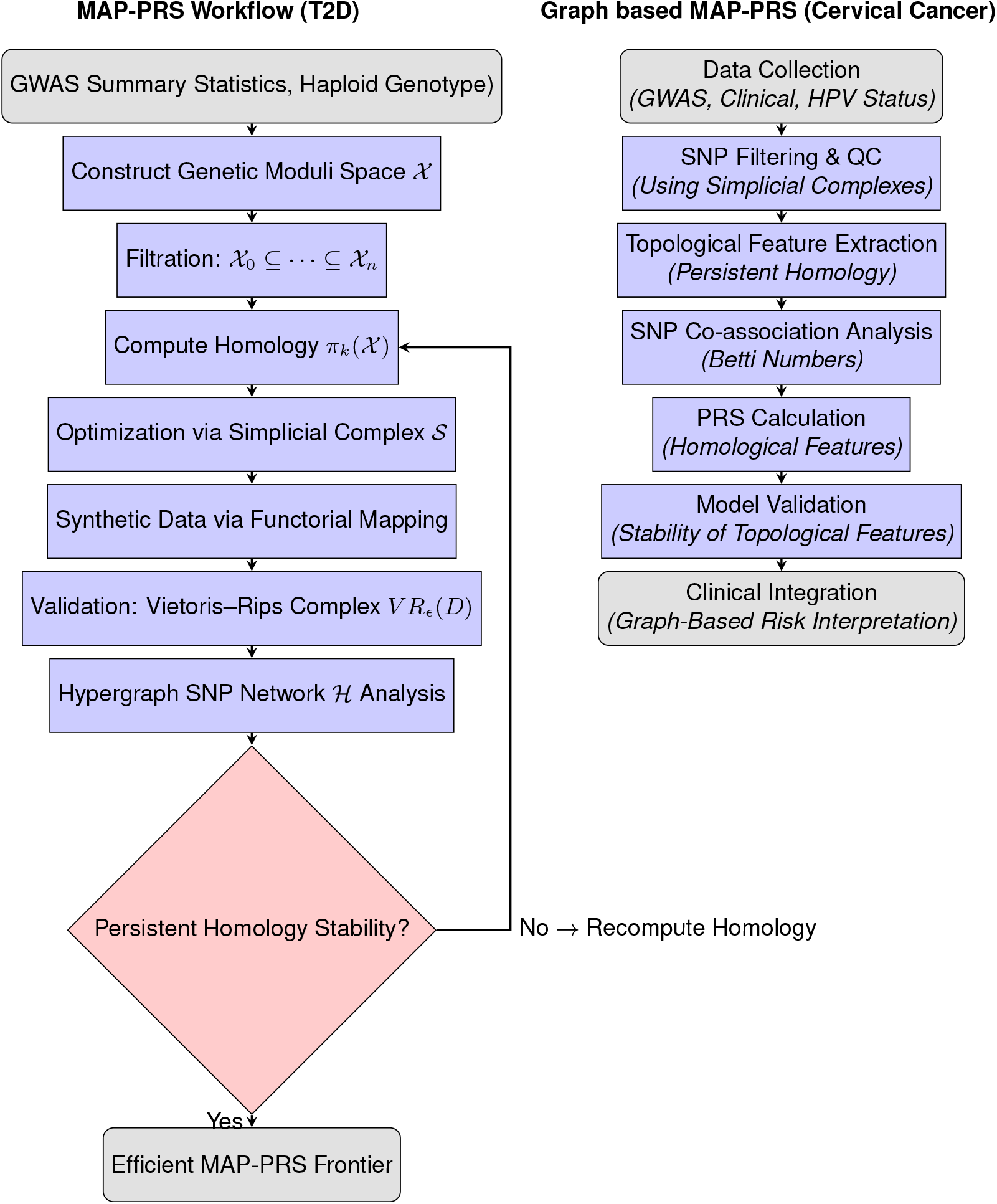
MAP-PRS Implementation

1. **GWAS summary statistics and Haploid genotype (input dataset)** step collect data from LDSC.^**9**^
2. **Preprocessing and SNP Filtering** step perform ancestry-aware QC and feature selection using simplicial complexes. Population-encoded data will allow for stratified filtering, ensuring that genetic variations relevant to different ancestries are appropriately accounted for.
3. **Model Development** step primarily aligns with the MAP-PRS workflow (left flowchart), where genetic data undergoes topological processing through the construction of a *genetic moduli space* (*χ*), *filtration*(*χ*_0_ ⊆ · · · ⊆ *χ* _*n*_), and *homology computation*(π_k_(*χ*)). These steps ensure that polygenic risk scores (PRS) are enriched with stable topological features. If large-scale GWAS summary statistics are un-available, case-control association studies will be conducted, which corresponds to SNP filtering and association analysis in the Graph-Based MAP-PRS for Cervical Cancer (right flowchart). **IV. Evaluation and Validation** step is reflected in both workflows but is more explicitly detailed in the MAP-PRS workflow, where *persistent homology stability* is assessed through *Vietoris–Rips complex validation* (*V R*_*ϵ*_(*D*)) and *hypergraph SNP network* (ℋ) analysis. The decision node in this workflow ensures that if homology stability is not met, recomputation occurs, ensuring model robustness. Meanwhile, the Graph-Based MAP-PRS (right flowchart) incorporates validation by testing the *stability of topological features* before clinical application. **V. Clinical Translation** step corresponds directly to the right flowchart, where PRS-derived risk predictions are integrated with real-world screening protocols. The use of *graph-based SNP networks and topological feature extraction* enables a more interpretable risk assessment, ultimately linking genetic insights to clinical decision-making.

## 7 Results

In this study, we implemented a European-Ancestry Portfolio-Based PRS approach using T2D GWAS to predict optimal efficient MAP-PRS frontier for T2D.

### 7.1 Persistent diagram

To integrate topological information into the MAP-PRS framework, we used *persistence diagrams* to summarize how groups of genetic variants (SNPs) connect and cluster together (Figure 10(a)). In simple terms, a persistence diagram shows which genetic connections appear, how long they last, and when they merge as we look across different genetic distance scales. Many short-lived *H*_0_ features near the diagonal represent SNP groups that are closely related and quickly merge together, reflecting the usual fine-scale linkage disequilibrium (LD) seen in genomic regions. In contrast, a few long-lived *H*_1_ features capture more persistent loops or cycles, indicating complex, higher-order relationships among SNPs that go beyond simple pairwise connections. These topological patterns provide a richer, multi-scale view of genetic structure. By combining this information with standard LD data, the MAP-PRS model can better recognize stable genetic patterns linked to disease risk, leading to more accurate and biologically meaningful risk predictions.

**Figure 10.**
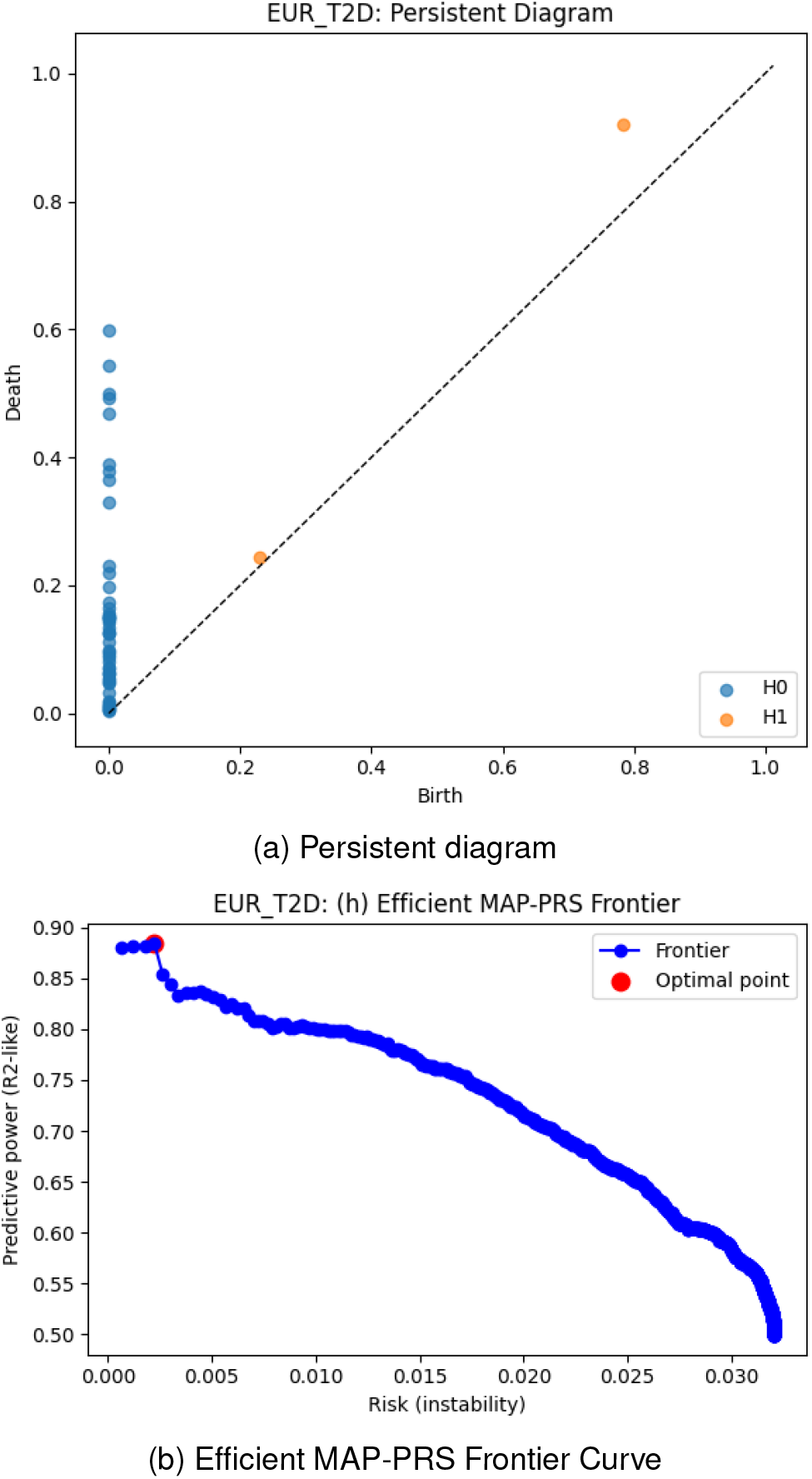
This visualization helps assess how well the PRS model predicts T2D risk in European ancestry.

### 7.2 Efficient MAP-PRS Frontier

The MAP-PRS framework leverages persistent homology for SNPs co-association analysis, effectively reducing the complexity of feature selection. As illustrated in Figure 10(b), the *efficient frontier optimization* within MAP-PRS identifies the optimal MAP-PRS point for type 2 diabetes (T2D), denoted by the red dot. This finding indicates that individuals of European ancestry exhibit robust optimal efficient MAP-PRS frontier because (i) good predictive performance, and (ii) its risk complexity represented by genetic uncertainty arising from linkage disequilibrium heterogeneity, allele-frequency divergence, and population instability is very low aligns with the studies.^**40, 41**^ Thus, establishing a foundation for broader, multi-ancestry implementation

This approach represents a promising step toward enabling its implementation in multi-ancestry settings, aiming to balance model generalizability and predictive power across diverse genetic backgrounds. By doing so, MAP-PRS ensures stable performance and improved robustness across populations with diverse ancestries.

## 8 Limitations and Future Work

### 8.1 Limitations

Obtaining the required sample size of the GWAS datasets, and haploid genotype for the ancestries is challenging.

### 8.2 Future Work

**Extension to Multi-Ancestry:** Building on our initial implementation, we are integrating additional components such as reinforcement learning-based SNP selection, efficient MAP-PRS frontier optimization, and computational performance analysis. While the current implementation has been evaluated using European-ancestry GWAS data for type 2 diabetes (T2D), we plan to extend this framework to include diverse ancestries and additional complex traits.

Ultimately, we aim to conduct clinical validation in collaboration with our clinical partners to evaluate the translational potential of our approach for real-world patient risk stratification across populations.

## Acknowledgements

We are grateful to Dr. Eyal Y. Kimchi (Department of Neurology, Northwestern University Feinberg School of Medicine, USA), Dr. Strajit Ghosh (Massachusetts General Hospital and Harvard Medical School, USA), Dr. Fernando Gómez-Baquero (Jacobs Technion–Cornell Institute, Cornell Tech; NSF Upstate New York Energy Storage Engine, USA), Dr. Russell Rockne, is an Associate Professor in the Department of Computational and Quantitative Medicine within Beckman Research Institute of City of Hope. He also serves as director of the Division of Mathematical Oncology, with the goal of translating mathematics, physics and evolution-based research to clinical care, Dr. Kamana Porwal (Department of Mathematics, IIT Delhi, India), and Dr. Mustafa Hajij (MSDSAI Program, University of San Francisco, California, USA) for their valuable guidance, support, and insightful suggestions, which greatly contributed to this work.

## Author Contributions

-Dr. Lokendra S. Thakur conceived the idea, designed the study, developed method and algorithm, curated data and done analysis, developed pipeline, all sections writing. - Dr. Nilanchali Singh performed the MAP-PRS clinical translation, and gynecological cancers related analysis. - Dr. Gurpreet Bharj performed the clinical robustness of the MAP-PRS. - All authors reviewed and approved the final version.

## Ethics Approval and Consent to Participate

Not applicable.

## Data Availability

We accessed raw data from the publicly available repositories as mentioned in the data description section. Code supporting this study are available on reasonable request.

## Disclaimer

Preprints are preliminary reports that have not been peer reviewed. They should not be regarded as conclusive, guide clinical practice, or be reported in news media as established information.

## References

[1] Florian Privé, Bjarni J. Vilhjálmsson, Hugues Aschard, and Michael G.B. Blum. Making the most of clumping and thresholding for polygenic scores. The American Journal of Human Genetics, 105(6):1213–1221, 2019.

[2] Alicia R. Martin, Masahiro Kanai, Yoichiro Kamatani, Yukinori Okada, Benjamin M. Neale, and Mark J. Daly. Current clinical use of polygenic scores will risk exacerbating health disparities. Nature Genetics, 51(4):584–591, 2019.

[3] Shaun Purcell, Benjamin Neale, Kathe Todd-Brown, Lori Thomas, Manuel A. R. Ferreira, David Bender, Julian Maller, Pamela Sklar, Paul I. W. de Bakker, Mark J. Daly, and Pak C. Sham. Plink: A tool set for whole-genome association and population-based linkage analyses. American Journal of Human Genetics, 81(3):559–575, 2007.

[4] Bjarni J. Vilhjálmsson, Jian Yang, Hilary K. Finucane, Alexander Gusev, Sara Lindström, Stephan Ripke, Giulio Genovese, Po-Ru Loh, Gaurav Bhatia, Ron Do, Tyler Hayeck, Hyejung Won, Sekar Kathiresan, Carlos Pato, Michele Pato, Nicholas Tam, Eli A. Stahl, Noah Zaitlen, Bogdan Pasaniuc, Gillian Belbin, Eimear E. Kenny, Philip L. de Jager, Nikolaos A. Patsopoulos, Steven McCarroll, Mark Daly, Shaun Purcell, Daniel I. Chasman, Benjamin M. Neale, Eric Lander, Peter M. Visscher, Peter Kraft, Nick Patterson, and Alkes L. Price. Modeling linkage disequilibrium increases accuracy of polygenic risk scores. American Journal of Human Genetics, 97(4):576–592, 2015.

[5] Tian Ge, Chia-Yen Chen, Guanghao Ni, Yun Feng, Jordan W. Smoller, and Xihong Lin. Polygenic prediction via bayesian regression and continuous shrinkage priors. Nature Communications, 10(1):1776, 2019.

[6] Xiang Zhou, Peter Carbonetto, and Matthew Stephens. Polygenic modeling with bayesian sparse linear mixed models. PLoS Genetics, 9(2):e1003264, 2013.

[7] Timothy S. H. Mak, Raphael M. Porsch, Seung Hoan Choi, Xiang Zhou, Pak C. Sham, and Yun S. Song. Polygenic scores via penalized regression on summary statistics. Genetic Epidemiology, 41(6):469–480, 2017.

[8] Alexandre Roth, Alicia R. Martin, Benjamin M. Neale, Simon Gravel, and Noah Zaitlen. Polygenic prediction across populations is influenced by ancestry, genetic architecture, and methodology. Cell Genomics, 3(10):100408, 2023.

[9] Hilary K. Finucane, Brendan Bulik-Sullivan, Alexander Gusev, Gosia Trynka, Yakir Reshef, Po-Ru Loh, Verneri Anttila, Han Xu, Chongzhi Zang, Kyle Farh, Stephan Ripke, Felix R. Day, Shaun Purcell, Eli A. Stahl, Sara Lindstrom, John R. B. Perry, Yukinori Okada, Soumya Raychaudhuri, Mark J. Daly, Nick Patterson, Benjamin M. Neale, and Alkes L. Price. Partitioning heritability by functional annotation using genome-wide association summary statistics. Nature Genetics, 47(11):1228–1235, 2015.

[10] Harry M. Markowitz. Portfolio selection. The Journal of Finance, 7(1):77–91, 1952.

[11] Stephen Boyd, Kristoffer Johansson, Richard Kahn, and et al. Markowitz portfolio construction at seventy. arXiv preprint, 2024.

[12] Andreas Dannenberg. Portfolio optimization under uncertainty. arXiv preprint, 2009.

[13] C. Bhattacharyya, K. Subramanian, B. Uppili, et al. Mapping genetic diversity with the genomeindia project. Nature Genetics, 57(4):767–773, 2025.

[14] Cathie Sudlow, John Gallacher, Naomi Allen, Valerie Beral, Paul Burton, John Danesh, Paul Downey, Paul Elliott, Jane Green, Martin Landray, Bette Liu, Paul Matthews, et al. Uk biobank: an open access resource for identifying the causes of a wide range of complex diseases of middle and old age. PLoS Medicine, 12(3):e1001779, 2015.

[15] Clare Bycroft, Colin Freeman, Desislava Petkova, Gavin Band, Lloyd T. Elliott, Kevin Sharp, Allan Motyer, Damjan Vukcevic, Olivier Delaneau, Jared O’Connell, Adrian Cortes, Samantha Welsh, Alan Young, Mark Effingham, et al. The uk biobank resource with deep phenotyping and genomic data. Nature, 562(7726):203–209, 2018.

[16] Genevieve L. Wojcik, Mariaelisa Graff, Kimberly K. Nishimura, Rui Tao, Jennifer Haessler, Christopher R. Gignoux, Heather M. Highland, Yesha M. Patel, Eric P. Sorokin, Crystal L. Avery, Gina M. Belbin, Stephanie A. Bien, Iona Cheng, Sarah Cullina, et al. Genetic analyses of diverse populations improves discovery for complex traits. Nature, 570(7762):514–518, 2019.

[17] John Michael Gaziano, John Concato, Mary Brophy, Louis Fiore, Saiju Pyarajan, James Breeling, Stacey Whitbourne, Jennifer Deen, Colleen Shannon, et al. Million veteran program: A mega-biobank to study genetic influences on health and disease. Journal of Clinical Epidemiology, 70:214–223, 2016.

[18] Akiko Nagai, Makoto Hirata, Yoichiro Kamatani, Kaori Muto, Koichi Matsuda, Yutaka Kiyohara, Toshiharu Ninomiya, Akiko Tamakoshi, Zentaro Yamagata, Taisei Mushiroda, Yoshinori Murakami, Koichiro Yuji, Yoichi Furukawa, Hitoshi Zembutsu, Toshihiro Tanaka, et al. Overview of the biobank japan project: Study design and profile. Journal of Epidemiology, 27(3S):S2–S8, 2017.

[19] Sudmant P., Rausch T., Gardner E., and others. An integrated map of structural variation in 2,504 human genomes. Nature, 526(7571):75–81, 2015.

[20] J. R. Munkres. Elements of Algebraic Topology. Addison-Wesley, 1984.

[21] A. Hatcher. Algebraic Topology. Cambridge University Press, 2002.

[22] J.-D. Boissonnat, F. Chazal, and M. Yvinec. Geometric and Topological Inference. Cambridge University Press, 2018.

[23] H. Edelsbrunner, D. Letscher, and A. Zomorodian. Topological persistence and simplification. Discrete & Computational Geometry, 28(4):511–533, 2002.

[24] A. Zomorodian and G. Carlsson. Computing persistent homology. Discrete & Computational Geometry, 33(2):249–274, 2005.

[25] J.-C. Hausmann. On the vietoris-rips complexes and a cohomology theory for metric spaces. In F. Quinn, editor, Prospects in Topology, pages 175–188. Princeton University Press, 1995.

[26] Yang Y, Tao R, Shu X, et al. Incorporating polygenic risk scores and nongenetic risk factors for breast cancer risk prediction among asian women. JAMA Network Open, 5(3):e2290174, 2022.

[27] Heather E. Wheeler, Keston Aquino-Michaels, et al. Poly-omic prediction of complex traits : Omickriging. arXiv preprint, arXiv:1303.1788, 2013.

[28] M. Shi, O’Brien, K.M., and C.R. Weinberg. Interactions between a polygenic risk score and non-genetic risk factors in young-onset breast cancer. Scientific Reports, 10(1):3242, 2020.

[29] Monica Isgut, Andrew Hornback, Yunan Luo, et al. Are gene-by-environment interactions leveraged in multi-modality neural networks for breast cancer prediction? arXiv preprint, arXiv:2407.20978, 2024.

[30] Ning Yu, Ke Li, Peng Zhou, Jitendra Malik, Larry S. Davis, and Mario Fritz. Inclusive GAN: Improving data and minority coverage in generative models. In Proceedings of the European Conference on Computer Vision (ECCV), 2020.

[31] Junjie Chen, Mohammad Erfan Mowlaei, and Xinghua Shi. Population-scale genomics data augmentation based on conditional generative adversarial networks. In Proceedings of the 11th ACM International Conference on Bioinformatics, Computational Biology and Health Informatics (BCB ‘20), pages 1–6, 2020.

[32] D. Machado Reyes, A. Bose, E. Karavani, and L. Parida. Fairprs: Adjusting for admixed populations in polygenic risk scores using invariant risk minimization. Pacific Symposium on Biocomputing, 28:198–208, 2023.

[33] Y. Shi, L. Li, Z. Hu, S. Li, S. Wang, J. Liu, Z. He, J. Zhou, J. Chen, C. Chen, Y. Liu, X. Zhang, N. Li, Y. Qiao,D. Ma, H. Shen, X. He, K. Zhai, and W. Lu. A genome-wide association study identifies two new cervical cancer susceptibility loci at 4q12 and 17q12. Nature Genetics, 45(8):918–922, 2013.

[34] D. Chen, I. Juko-Pecirep, J. Hammer, E. Ivansson, S. Enroth, I. Gustavsson, R. S. Fehrmann, T. Torngren,C. Williams, S. Holm, B. Andersson, J. Zubovits, M. Tommasino, J. Dillner, and U. Gyllensten. Genome-wide association study of susceptibility loci for cervical cancer. Journal of the National Cancer Institute, 105(9):624–633, 2013.

[35] Anika Misra, Buu Truong, Sarah M. Urbut, Yang Sui, Akl C. Fahed, Jordan W. Smoller, Aniruddh P. Patel, and Pradeep Natarajan. Instability of high polygenic risk classification and mitigation by integrative scoring. Nature Communications, 16(1):1584, 2025.

[36] Saunders Mac Lane. Categories for the Working Mathematician. Springer, 1971.

[37] Steve Awodey. Category Theory. Ebsco Publishing, 2006.

[38] Brendan Fong and David I. Spivak. An Invitation to Applied Category Theory: Seven Sketches in Compositionality. Cambridge University Press, 2019.

[39] Benjamin C. Pierce. Basic Category Theory for Computer Scientists. MIT Press, 1991.

[40] Moreno-Grau S., Vernekar M., Lopez-Pineda A., et al. Polygenic risk score portability for common diseases across genetically diverse populations. Human Genomics, 18(1):93, 2024.

[41] L. Duncan, H. Shen, B. Gelaye, et al. Analysis of polygenic risk score usage and performance in diverse human populations. Nature Communications, 10(1):3328, 2019.

